# Modeling Uncertainty in Grapevine Powdery Mildew Epidemiology Using Fuzzy Logic

**DOI:** 10.1101/264622

**Authors:** R. A. Choudhury, W. F. Mahaffee, N. McRoberts, W. D. Gubler

**Author notes:** Corresponding author: W. D. Gubler. Mention of trade names or commercial products is solely for the purpose of providing specific information and does not imply recommendation or endorsement by the U. S. Department of Agriculture. USDA is an equal opportunity provider and employer.

## Abstract

Powdery mildew is the most important disease of grapevines worldwide. Despite the potential for rapid spread by the causal pathogen, grape powdery mildew has been effectively managed using fungicide applications applied based on a calendar schedule or modeled disease risk index. Various epidemiological models for predicting disease development or risk have helped to improve disease management. The Gubler-Thomas (GT) risk index is a popular disease risk model used by many growers in the western U.S. We modified the GT risk index using fuzzy logic to address both biological and mechanical uncertainty in the pathosystem. The spraying schedule suggested by the fuzzy-modified GT risk index was tested in eight site-years. Overall, the fuzzy-modified risk index maintained comparable levels of disease control as both the original model and a calendar based treatment, and had significantly less disease than the untreated control. The fungicide use efficiency of the fuzzy-modified GT risk index suggests that the updated risk index was significantly more efficient with fungicide applications than both the calendar and original GT risk index.

## INTRODUCTION

Grapevine powdery mildew, caused by *Erysiphe necator* (Schwein.), affects all cultivated *Vitis vinifera* grapevines, and is the most serious persistent threat to grape production world. Powdery mildew epidemics typically begin in the early spring with the release of ascospores from overwintering ascocarps lodged in the bark of grapes during periods of wetness (Gadoury and Pearson 1988). Once the ascospores have infected green host tissue, the pathogen begins to reproduce asexually. During periods of conducive weather conditions, rapid proliferation of asexual conidia on susceptible tissue can lead to extremely damaging epidemics if the pathogen is left unchecked.

Many California and Oregon grape growers apply fungicides on a calendar schedule based on the shortest recommended application interval. However, this can often lead to over-application of fungicides if disease pressure is low or if the host is relatively resistant. Several epidemiological models have been created to predict disease risk in an effort to reduce and optimize fungicide use (Chellemi and Marois 1991; Gubler et al. 1999; Sall 1980). These epidemiological models differ in their data input as well as their ability to predict future disease risk. The most widely adopted epidemiological model in California is the Powdery Mildew Risk Assessment Index, also known as the Gubler-Thomas (GT) risk index (Gubler et al. 1999; Lybbert et al. 2016; Lybbert and Gubler 2008; Thomas et al. 2002).

The GT risk index uses temperature and leaf wetness to first predict ascospore release and initiates a risk index. The risk index ranges from 0 to 100, and functions as an advisory tool that growers can use to efficiently choose the fungicide application interval or chemistry. The GT risk index was shown to maintain disease control while eliminating from 2 to 8 fungicide applications compared with a calendar-based schedule (Gubler and Thomas 2006) and used by over 50% of surveyed grape growers and managers (Lybbert and Gubler 2008). Despite its successes, the GT risk index tends to overestimate risk which results in excess applications of fungicide (Lybbert et al. 2016). However, the conservative approach has prevented management failures associated with use of the GT risk index for nearly 20 years.

Recent research has examined how weather conditions impact the ability of *E. necator* to grow and reproduce (Choudhury et al. 2014; Moyer et al. 2010; Peduto et al. 2013). Peduto et al. (2013) effectively combined a series of controlled environment studies and field trials to demonstrate that improvements to the GT risk index are possible. Other recent work using controlled environment studies has found that *E. necator* can withstand repeated exposure to temperatures that were thought to be lethal or sub-lethal (Choudhury et al. 2014). The success of the Peduto et al. (2013)-altered GT index with a higher lethal temperature threshold suggests that *E. necator* is tolerant of excessive heat in both field and laboratory settings. Many of these controlled environment studies suggest that temperature drives most of the life-history of the pathogen. However, other important confounding environmental factors such as relative humidity, ultraviolet radiation, and free moisture can also play a large role in its growth and reproduction (Austin and Wilcox 2012; Carroll and Wilcox 2003; Vergely 2002). Since the extent and form of the uncertainty in pathogen response to these variables are themselves unknown, it is difficult to account for them using parametric approaches (Scherm 2000). One way to capture such uncharacterized uncertainty in dynamic processes is to use fuzzy logic (Zadeh 1965).

While classical logic is based on assigning binary “true” or “false” truth values, many biological concepts are inherently uncertain because they may not be easily defined in terms of binary outcomes. While there may be variation within a single population, it is often possible to assign membership values which describe the extent to which individuals can be considered to belong to different categories (Zimmermann 2001). In fuzzy logic, values that have no uncertainty are considered “crisp”. Values that are “fuzzy” have uncertainty associated with them and this uncertainty can be visualized and expressed as a polygon around the “crisp” value in a 2-dimensional plane. The abcissa of the plane is the certainty function with a minimum of zero and a maximum of one, while the ordinate measures the possible values of the fuzzy quantity. In this way, crisp numbers are simply a special case of fuzzy numbers in which all the certainty is concentrated on a single value, and the polygon collapses to vertical line of height 1.

The certainty (or belief) function for a fuzzy number can also be considered as a set membership function (Zimmermann 2001). As a consequence of this homology between fuzzy numbers and sets, the overlap between two fuzzy numbers can be considered as a fuzzy set intersection (Zimmermann 2001). Such intersections can be used as a decision tool in disease forecasting (Kim et al. 2005), summarizing the extent to which, for example, a measured temperature intersects with the range of suitable and unsuitable temperatures for pathogen growth.

The increasing popularity and use of personalized weather stations in precision agriculture has allowed many growers to implement site-specific disease forecast models (de Wolf and Isard 2007). There is, however, uncertainty associated with sensor accuracy, microclimates and variation (in some regions) in altitude that all contribute to a significant amount of variation in measured values from true values (Pfender et al. 2012).

In many epidemiological models weather data are processed as crisp numbers regardless of whether they are derived from averages over a period of time or discrete measurements at a specific time. Both approaches generally disregard the variance associated with measurement itself. However, different classes of weather stations that vary in their accuracy depending on the parameter being measured and sensor used (Pfender et al. 2011). These uncertainties are further complicated by the uncertainties in estimations of the optimal and lethal temperatures for pathogen growth based on controlled laboratory studies (Chellemi and Marois 1991; Delp 1954; Peduto et al. 2013) where sensor error and spatial variability of the experimental parameter being manipulated within the chamber.

In this study, we modified the existing GT risk index for grape powdery mildew using fuzzy logic to account for uncertainty in both weather and biological data. We then tested fungicide spray schedules suggested by the original GT index and the modified fuzzy GT index in eight test vineyards, and measured disease severity on both leaves and clusters at regular intervals throughout the season. This allowed us to directly compare the effectiveness of the original GT index and the fuzzy GT index. The original GT risk index was used as a benchmark for comparison on two criteria: reduction in the number of fungicide applications and efficacy of disease control.

## MATERIALS AND METHODS

### Creation of the fuzzy GT risk index

Fuzzy logic was used to modify the Gubler-Thomas (GT) powdery mildew risk index model (Gubler et al. 1999). The model consists of several if-then statements that help guide whether the index is active or not. If there have been 3 consecutive days with at least 6 hours of optimal conditions post budbreak, then trigger the index; within each 24 hour period if there have been 6 hours of optimal conditions, then add 20 points to the index; if there have not been at least 6 hours of optimal conditions, then subtract 20 points from the index; if there have been temperatures greater than 35 °C for at least 15 minutes, then subtract 10 points from the index. The index cannot increase more than 20 points or decrease more than 10 points in a single day, and is constrained to a maximum value of 100 and a minimum of 0.

The main aim of the modifications made in this study was to deal with uncertainty in the way the GT risk index accounts for the effects of high temperature on pathogen development. By assuming a crisp threshold between suitable and unsuitable ambient temperatures the original GT risk index does not allow for high temperature adaptability in the pathogen. Model parameters of the original index were based on field and laboratory observations of optimal and lethal conditions of powdery mildew germination, growth, and sporulation (Delp 1954; Gubler et al. 1999).

To adapt the GT index the original risk index was first analyzed as a set of membership functions (Fig. 1). The crisp membership functions of the original model were then modified to the produce the fuzzy GT model using a combination experimental data and expert knowledge (Fig. 1). Estimation of uncertainty in weather station data was based on known variability in weather station data (Pfender et al. 2011). High accuracy for temperature is considered to be +/– 0.3°C whereas +/–1°C is a more commonly obtained range in practice and even this variability may be exceeded if sensors are not properly maintained and recalibrated. Averaging across a time interval (e.g. 1 min) can reduce the effect sensor inaccuracy but the more common practice of averaging across longer time intervals (e.g. 15 min or 1 hour) introduces the variance of the change in temperature over time due to air turbulence and mass transport into the uncertainty (Bailey et al. 2014).

**Fig. 1.**
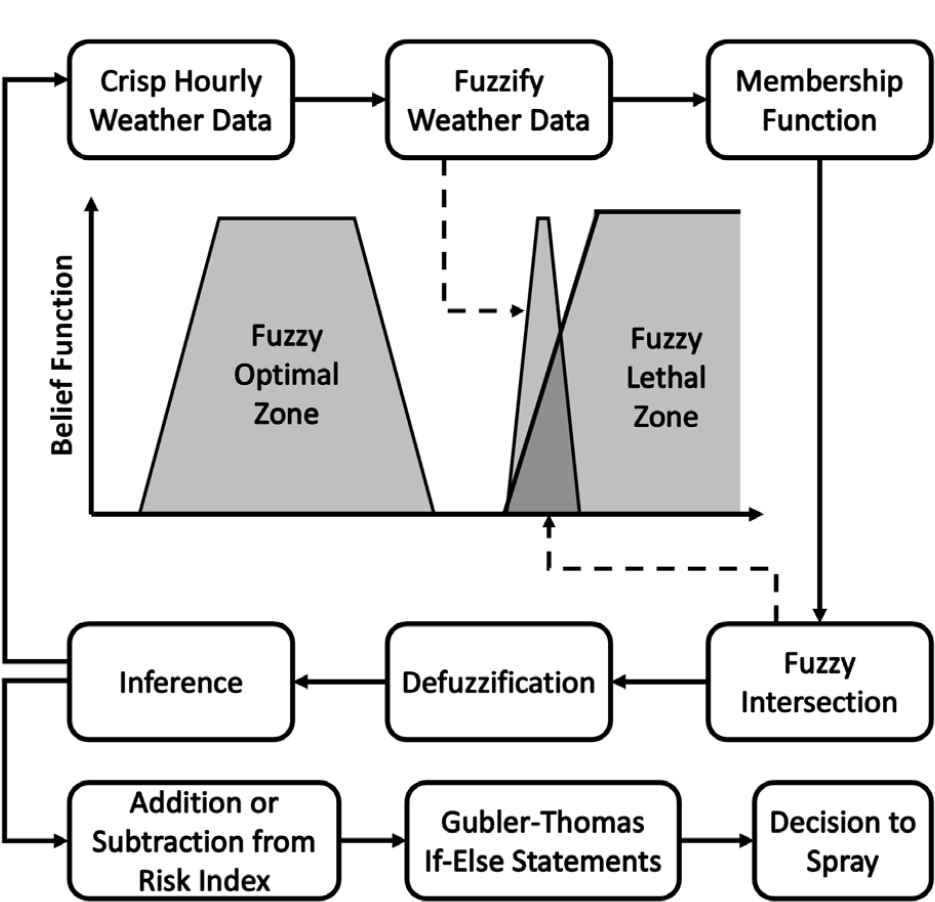
Cartoon schematic diagram of the fuzzy risk index model.

Estimations for the optimal and lethal growth ranges were based on controlled environment studies (Austin and Wilcox 2012; Delp 1954; Peduto et al. 2013). The upper threshold of the optimal growth range was reduced from the original GT risk index to account for deleterious effects of exposures to sub-lethal temperatures (Peduto et al. 2013). This change was supported by work that suggested that stress from other environmental factors can exacerbate the detrimental effects of increased heat (Austin and Wilcox 2012; Delp 1954). Estimation of the lethal growth range was based on controlled environment studies that suggest that the pathogen can be detrimentally affected by duration exposure of high temperatures, depending on other ambient environmental conditions (Austin and Wilcox 2012; Delp 1954; Peduto et al. 2013).

### Weather data

For field testing, temperature and rainfall data were imported from the weather station located closest to the field site. The Davis, Clarksburg, Corvallis, and Fresno site-years had an in-vineyard weather station (iMETOS ag, Pessl Instruments, GmbH) with hourly weather data that was downloaded daily. The Napa site-years relied on public weather stations ranging from 3 to 7 km from the test vineyard. After the weather data was imported into the software, hourly temperature data were adjusted according to the fuzzy membership function (Fig. 1). The fuzzy temperature data were then overlaid on top of the fuzzy membership functions to assess the intersection with these sets (Fig. 1). The fuzzy intersection area was calculated hourly using integration methods, and was then used as a decision criterion in the calculation of the fuzzy risk index value. For each hourly average temperature measurement If the area of the fuzzy intersection between the measured temperature and the lethal high temperature threshold, or optimal temperature range was greater than 0.6, then the hour was considered lethal/optimal or lethal for fungal growth, depending on which intersection was relevant for that hour.

### Weather data

For field testing, temperature and rainfall data were imported from the weather station located closest to the field site. The Davis, Clarksburg, Corvallis, and Fresno site-years had an in-vineyard weather station (iMETOS ag, Pessl Instruments, GmbH) with hourly weather data that was downloaded daily. The Napa site-years relied on public weather stations ranging from 3 to 7 km from the test vineyard. Weather stations export hourly data as crisp numbers. After the weather data was imported into the software, hourly temperature data were fuzzified according to the fuzzy membership function. The fuzzy temperature data were then overlaid on top of the fuzzy optimal and fuzzy lethal membership functions to assess the fuzzy intersection with these fuzzy sets; i.e. the area of overlap between membership functions for the temperature value and each of the growth-response functions. The fuzzy intersection values were used as a decision tool. If the area of the fuzzy intersection was greater than 0.6, then the hour was considered optimal or lethal for fungal growth, depending on the comparison in question. The threshold fuzzy intersection value of 0.6 was determined empirically based on previous weather data and powdery mildew disease severity.

### Field validation

Randomized complete block design experiments were implemented in eight vineyards over two years to compare the fungicide schedule suggested by the GT and fuzzy GT risk indices. Field sites in Clarksburg and Napa planted with ‘Chardonnay’ grapes; those in Davis and Fresno were planted with ‘Thompson Seedless’; and the Corvallis site was planted with ‘Pinot Noir’. In California, each treatment replicate consisted of three vines, and each replicate was repeated six times in a randomized complete block experiment. Field sites in Davis, Clarksburg, and Fresno included an untreated control treatment as well as a fuzzy GT, GT, and calendar spray programs. An untreated control treatment was not feasible in either of the Napa sites due to potential financial impacts on neighboring vineyards arising from the presence of untreated blocks.

For disease control three different fungicides were applied in rotation: Quintec (quinoxyfen; Dow AgroSciences LLC), Luna Experience (fluopyram + tebuconazole; Bayer CropScience LP), and Flint (trifloxystrobin; Bayer CropScience LP) in 2011; and Quintec, Adament (tebuconazole + trifloxystrobin; Bayer CropScience LP), and Flint in 2012. The fungicide regime was changed between 2011 and 2012 due to concern about local populations of the pathogen developing fungicide resistance. All fungicides were used at their recommended application rates, and were applied on 14-21 day intervals, depending on the values of the risk indices. Fungicide applications and leaf disease severity ratings began when shoots reached a length of 30cm height, following common grower practices in California. The calendar spray program used the three different fungicides in rotation, applied on a 14 day interval.

Eighteen leaves from each replicate were assessed weekly or biweekly for disease severity, a percentage estimate of the leaf area covered with mildew. Six leaves from each vine were randomly selected from the fruit zone. Disease ratings always occurred before fungicide applications to ensure safety for the disease raters. When grape berries were pea-sized, eighteen clusters per replicate were rated for severity in addition to leaves. Grape cluster severity is a percentage estimate of the amount of visible cluster surface covered with powdery mildew mycelia. Leaves were removed from the canopy for disease rating while clusters were left intact on the vine. In the field, ratings were randomly performed by several disease raters. Disease ratings and fungicide applications continued until grape berries reached 10° Brix, following common grower practices (Gadoury et al. 2003; Gubler et al. 1999).

The field site in Corvallis, OR also used a randomized complete block design, with six blocks and each replicate consisting of five vines. The trial consisted of four treatments: a GT index treatment, a fuzzy GT index treatment, a calendar treatment, and an untreated control, which was sprayed with water at the same time as the GT treatment. Disease incidence was measured weekly and severity ratings on leaves were measured monthly. For disease incidence, 10 leaves on the 6^th^ or 7^th^ node from the growing tip for each were rated as disease if single colony was observed on the adaxial or abaxial surface. Each month disease severity was assessed from the same leaves and recorded as an average of percent leaf area covered on the abaxial and adaxial sides of the leaf. At véraison, ten clusters per vine from the middle three vines were harvested and frozen at −20°C. Twenty-five berries per cluster were arbitrarily selected and microscopically examined for presence of *E. necator* infection. Two blocks were rated for disease severity for each of the treatments using a five-category ordinal scale. Disease severity ratings were then adjusted to represent the percent of the maximum score.

### Data analysis

Disease severity on both clusters and leaves was analyzed for the entire season for all site years. True means and standard error of the means were calculated for all sites. In addition to true means, disease severity on both clusters and leaves was analyzed using a linear mixed model approach for all sites-years using the R package lme4 (Bates et al. 2014). The mixed model consisted of fungicide scheduling treatments as a fixed effect and block, site, and year as random effects. All site-years had a GT treatment and a fuzzy GT treatment. All site-years except those conducted in Napa, CA were conducted with an untreated control as well. Pairwise least square means comparisons between treatments were conducted using a Tukey’s honestly significant difference (HSD) test adjustment for all site-years except those conducted in Napa, CA. In the Napa, CA site-years, means comparisons were conducted using Student’s t test, because there were only two treatments. Risk indices were analyzed for the average risk index and the area under the curve of the index.

Area under the disease progress curve (AUDPC) for both cluster and leaf disease severity data was calculated using the audpc() function in the agricolae package in R (De Mendiburu 2014). AUDPC was calculated at the block level for all site-years. Fungicide use efficiency (*E*) was calculated for the calendar, fuzzy logic, and GT treatments following the modified equation used of Small et al. (2015). Fungicide use efficiency is a measure of the percent of disease control per fungicide application, and is caluculated as:

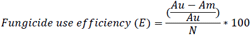

in which *Au* is the AUDPC for the untreated crop, *Am* is the AUDPC for a spray schedule method, and *N* is the number of fungicide applications scheduled by the method. AUDPC and fungicide use efficiency values were compared using a linear mixed model approach as described above.

## RESULTS

### Implementation of the fuzzy model

Fuzzy modeling was first implemented in 2011 using FuziCalc software (FuziWare Inc, Knoxville, TN) and then using R v. 2.10.0 (www.cran-project.org) and Excel 2007 (Microsoft Corporation, Redmond, WA) in 2012. The software was changed from FuziCalc to R/Excel because of the wider availability of R and Excel. While both software packages produced the same output, they functioned in slightly different ways. FuziCalc has a graphical user interface, allowing users to define the specific shapes and coordinates of different membership functions. Weather data was first imported and then fuzzified to fit the membership function. A series of ‘if-then-else’ statements were used to decide whether temperature data was optimal or lethal (Fig. 1), and ultimately decide whether to increase or decrease the risk index. The use of FuziCalc was discontinued in this study because the software is no longer commercially available, and because the software would often not function on computers using modern operating systems. The R/ Excel software was processed by first fitting the weather data to the membership function in Excel, and then exporting to R to assess whether the fuzzy weather data overlaps with the optimal and lethal membership functions. Membership functions were represented as polygons in R. After assessing the fuzzy intersection in R, data was exported back to Excel. A series of ‘if-then-else’ statements were used to decide whether the fuzzy hourly weather data was optimal or lethal, and ultimately whether to add or subtract points from the risk index. After creating the R/ Excel fuzzy model, the same weather data was used in both R/ Excel and FuziCalc to ensure that the R/ Excel model created the same risk index as the FuziCalc model.

### Weather data

Weather stations export hourly data as crisp numbers. After the weather data was imported into the software, hourly temperature data were fuzzified according to the fuzzy membership function. The fuzzy temperature data were then overlaid on top of the fuzzy optimal and fuzzy lethal membership functions to assess the fuzzy intersection with these fuzzy sets; i.e. the area of overlap between membership functions for the temperature value and each of the growth-response functions. The fuzzy intersection values were used as a decision tool. If the area of the fuzzy intersection was greater than 0.6, then the hour was considered optimal or lethal for fungal growth, depending on the comparison in question. The threshold fuzzy intersection value of 0.6 was determined empirically based on previous weather data and powdery mildew disease severity.

### Field validation

Field trials comparing the fuzzy GT and original GT risk indices were conducted at eight site-years. Disease developed early on both untreated leaves and clusters at the Clarksburg site-years. Disease severity at the Davis and Fresno site-years were lower than those at Clarksburg, possibly due to differential resistance of varieties. Although little to no disease developed at the Napa site-years, vineyards in the surrounding area were affected by powdery mildew, suggesting the presence of regional inoculum (R.A. Choudhury, personal observation).

### Risk indices

The two risk indices differed in all eight site-years (Fig. 2). The mean risk index for the fuzzy GT risk index across all site-years was 14 points lower than the original GT risk index (Fig. 2). This difference between the models was especially noticeable in warmer areas (e.g. - Fresno) where the fuzzy model predicted lower disease pressure than the original model as the season progressed. Although there were broad differences between the two indices across the study, the mean risk indices did not significantly differ at the Corvallis-2012, Napa-2011, or Napa-2012 site-years. The fuzzy GT risk index had on average slightly fewer fungicide applications and a longer application interval (Fig. 2). The fuzzy GT risk index was able to reduce fungicide applications while maintaining comparable disease control to the original GT risk index at four of the eight site-years (Fig. 2).

**Fig. 2:**
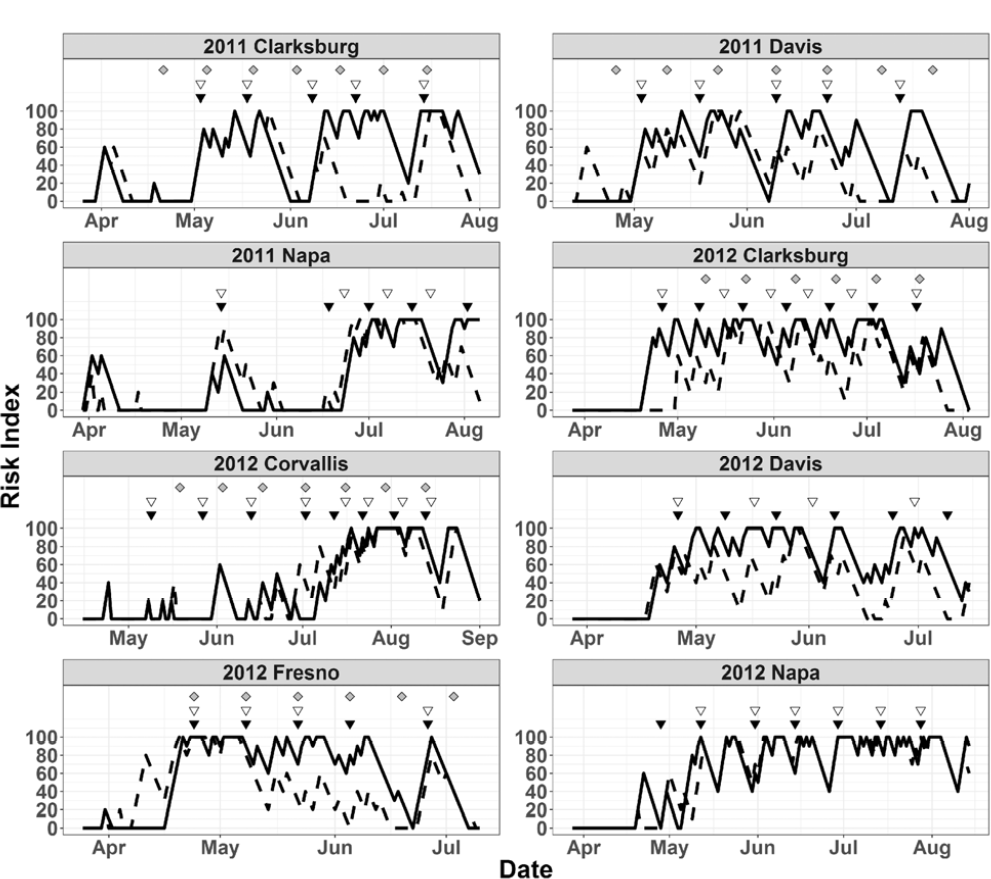
Grapevine powdery mildew risk indices and fungicide spray applications for the site-years in this study. Dashed line represents fuzzy GT index, solid line is the original GT index. Solid inverted triangles, empty triangles, and grey diamonds represent fungicide applications for the GT, fuzzy GT, and calendar treatments, respectively.

### Final disease severity following different indices

The mixed model analysis revealed significant differences in final disease severity in leaves and clusters at different site-years (Fig. 3). In general these significant effects could be attributed to the difference between the untreated control and the other treatments in site-years where the untreated control was present. There were no differences in disease leaf severity between the two index treatments at any of the site-years. Similar patterns of results were observed for cluster disease ratings. The untreated control treatments had consistently higher cluster disease severity at all site years. There were no statistical differences between the GT and fuzzy GT treatments in cluster disease severity at any of the site-years.

**Fig. 3:**
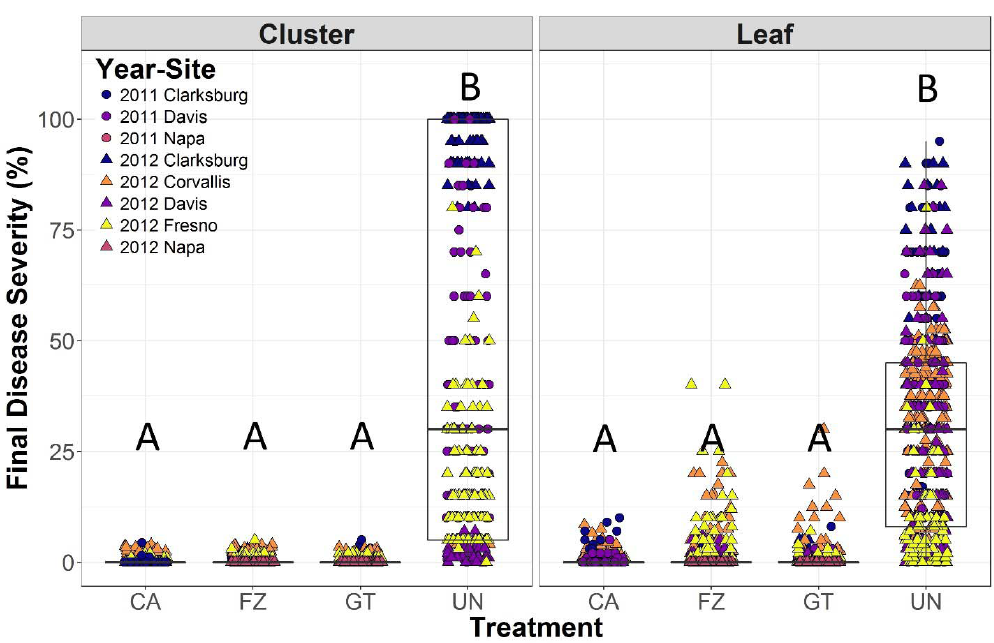
Final powdery mildew disease severity ratings for the four treatments: calendar (CA), fuzzy modified GT risk index (FZ), original GT risk index (GT), and untreated (UN). Lettering describes significant differences based on Tukey’s honestly significant difference (HSD) test at α=0.05.

### AUDPC and Fungicide use efficiency

The mixed model revealed significant differences in AUDPC in leaves and clusters at different site-years (Fig. 4). The mixed model analyses also revealed statistical differences in AUDPC where the untreated control had a consistently and significantly higher AUDPC than either of the index treatments or the calendar treatment at all site-years. However, there were no differences in AUDPC between the two index treatments or the calendar treatment at any of the site-years. The mixed model analyses also revealed statistical differences in fungicide use efficiency (Fig. 5). The fuzzy GT treatment control had a consistently and significantly higher fungicide use efficiency than either the original GT index treatment or the calendar treatment at all site-years tested whether considering both leaf or cluster disease data.

**Fig. 4:**
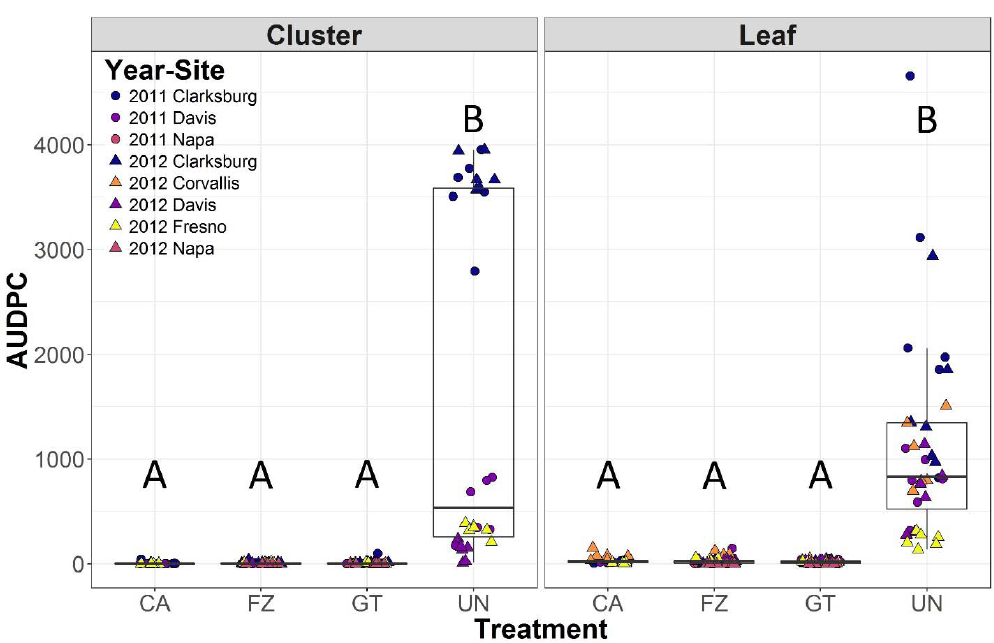
Area under the disease progress curve (AUDPC) for the four treatments: calendar (CA), fuzzy modified GT risk index (FZ), and original GT risk index (GT). Lettering describes significant differences based on Tukey’s honestly significant difference (HSD) test at α=0.05.

**Fig. 5:**
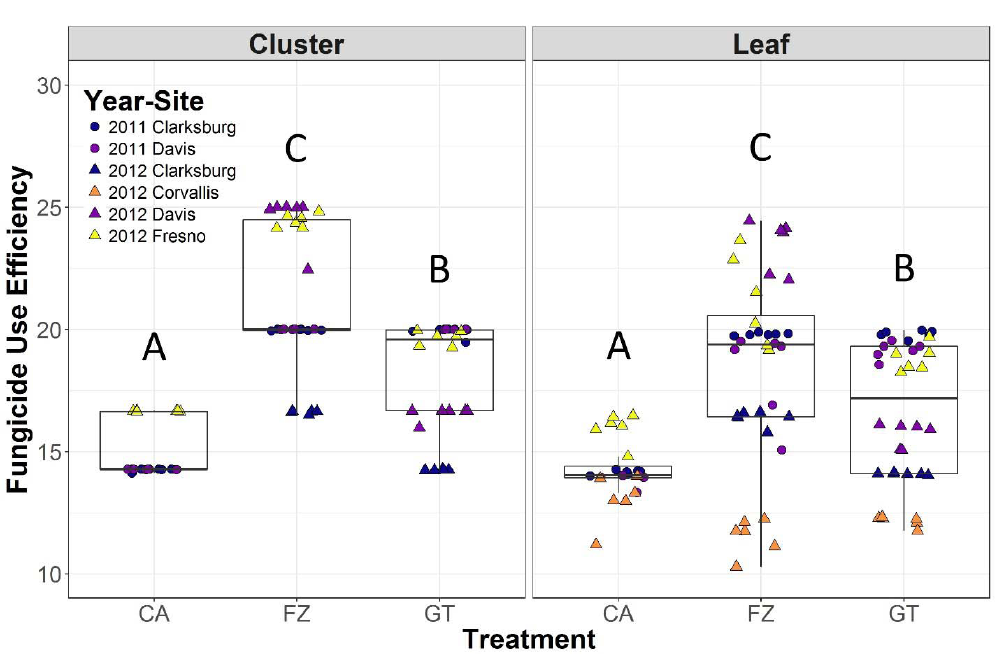
Fungicide use efficiency for the three spray schedules: calendar (CA), fuzzy modified GT risk index (FZ), original GT risk index (GT), and untreated (UN). Lettering describes significant differences based on Tukey’s honestly significant difference (HSD) test at α=0.05.

## DISCUSSION

The goals of this study were to modify an existing decision support system to address instrumental and biological uncertainty, and assess whether the updated model improved disease control and it impact on fungicide use. The fuzzy-modified GT risk index performed well compared with fungicide schedules suggested by the original GT risk index and a calendar-based spray schedule, reducing the number of fungicide applications at several sites without significant increases in disease severity. These data suggest that the fuzzy-modified GT risk index used fungicides more efficiently than the original GT risk index or a calendar-based spray schedule under the conditions tested.

Addressing sources of uncertainty in epidemiological models is one of the most important challenges for future plant disease research (Cunniffe et al. 2015). With the increasing availability of high resolution weather data as well as rapid and high throughput molecular diagnostics of local host populations (Thiessen et al. 2016), growers have more information than ever for assessing and responding to risk of disease. The availability of enabling technologies coupled with increasing interest and support for precision agriculture will only increase the importance of decision support systems in crop management (Gent et al. 2013; Shtienberg 2013). Ideally, uncertainty in weather data and the effects of that uncertainty on the accuracy of epidemiological models would be fully characterized and captured in appropriate probability distributions around model parameters. However, in practice models are often developed and deployed without such characterization and with little chance of the detailed work needed to achieve the results ever being carried out. In such situations an alternative approach to introducing uncertainty into existing models is needed, and the use of fuzzy arithmetic is one possible means to achieve that end.

While the fuzzy-modified GT risk index adjusted how the original GT risk index interpreted weather input variables and pathogen growth variables, it did not directly modify how the model interpreted time. A fuzzy modification of time could be very valuable, allowing for more flexibility in what conditions may be considered optimal or lethal (Choudhury et al. 2014; Peduto et al. 2013). In the original GT risk index, six consecutive hours of optimal temperatures are required to increase the risk index (Gubler et al. 1999). However, in the western USA where the GT model has been most widely adopted, during the warmer periods of the mid-season, it is common to have long periods of optimal temperature interspersed with mildly sub-optimal temperatures. While a single day may have more than six hours of optimal temperature, the interruptions to the optimal conditions mean that no increase the risk index occurs. A fuzzy interpretation of time requirements for pathogen development may help by recognizing these periods as optimal for growth of the pathogen.

Using fungicides efficiently is important, because it helps to reduce the economic and environmental effects of pesticides. There are over three hundred thousand hectares of grapes in California (USDA NASS 2016), and they are sprayed with approximately 8.1 million kilograms of sulfur (CDPR 2015), mostly for prophylactic treatment against powdery mildew. There have also been increases in uses of synthetic pesticides in grape production (CDPR 2015). Calculating the efficiency of fungicide use helps to elucidate how different DSS-scheduled sprays compare with one another (Small et al. 2015). Our study suggests that both the fuzzy modified GT risk index and the original GT risk index are more efficient in fungicide use than a calendar-based schedule of treatments.

Many growers rely on calendar-based pesticide applications as a form of crop insurance (Horowitz and Lichtenberg 1994). This can sometimes lead to perverse incentives when deciding whether to apply a pesticide or not, as growers frequently rely on pesticides more often than they need. The monetary costs associated with extra pesticide sprays (equipment use, labor, material) are often out-weighed by yield or quality losses, especially in high-value crops, or for diseases that are capable of rapid dispersal and require preventative treatment to maintain adequate control.

The GT model was originally developed with the aim of allowing growers to extend the interval between fungicide applications, when the risk of pathogen development is low. While the GT model has been adopted by many growers (Lybbert and Gubler 2008), they will frequently use the system to decide *what* to spray rather than *when* to spray, applying sulfur treatments when the risk index predicts low disease risk and synthetic pesticides under high disease risk (Lybbert et al. 2016). This alternative use of the original GT risk index may lead (counterintuitively) to an overall increase in the number of pesticides applied to the crop.

While several epidemiological models have been developed using fuzzy logic independent of existing models (Kim et al. 2005; Scherm 2000), we designed our fuzzy-modified GT risk index using the original GT risk index as a scaffold. Orlandini et al. (2003) similarly modified their PLASMO model using fuzzy logic, improving the model’s ability to predict grapevine downy mildew infection and disease intensity. Gonzalez-Dominguez et al. (2015) were also able to use fuzzy logic to match expert recommendations on grapevine downy mildew spray scheduling based on host and pathogen variables. Using the original GT risk index as a scaffold has the advantage of maintaining familiarity for growers (Lybbert and Gubler 2008) as well as a large retaining the relevance of the body of supporting literature and research. In a study of winegrape growers in the Lodi area of California, Hoffman et al. (2014) found that growers' perceptions of the financial value of a technology was correlated with their familiarity with the technology. The fuzzy-modified GT risk index maintains many of the features that growers valued from the original GT risk index and reduce the overall number of fungicides and maintain disease control in our eight site-years, suggesting the modified GT index, may have characteristics that would promote its adoption.

While modifying an existing decision support system has the advantage of familiarity to growers and an established infrastructure, there are some drawbacks. Modifying an existing decision support system restricts design capabilities compared with completely new systems. We maintained many of the attributes of the original GT risk index (e.g. - index minimum and maximum, points added and subtracted per day) (Gubler et al. 1999).

We developed the fuzzy modified risk index and then tested it in several sites over two years. We debated whether to update the fuzzy modified risk index after testing in the field for the first year in order to address how best to implement the fuzzy risk index. Dynamically adapting a decision support system as new data is collected would allow for a heuristic optimization of the model over the course of the study. However, these changes to the model would have reduced the number of repetitions of the direct comparison between treatments extending the development time, as each iteration would need to be independently tested at multiple sites and years to allow collection of sufficiently robust data. This dynamic creates an unusual tradeoff between creating an optimized system for the end user or presenting a model for publication in a peer-reviewed journal. Ideally, developers of a new DSS will be able to draw upon a large body of data before field testing (Gent et al. 2013; Shtienberg 2013). However, this is not conducive towards development of DSS’es for understudied pathosystems, or systems that experience different environmental conditions due to global climate change (Garrett et al. 2013). Changes in publication types and institutional rewards for developing tools may help to incentivize optimization techniques in DSS development (Howison and Herbsleb 2013).

There is uncertainty associated with every disease system. While controlled environment studies can help us understand important epidemiological factors that drive disease, there will always be confounding environmental factors that impact the host or pathogen in undescribed ways. Fuzzy logic is an effective way to address this uncertainty in plant disease epidemiology. The fuzzy GT risk index is a successful adaptation of the original GT risk index, and field testing at multiple sites in two years suggests that the fuzzy GT risk index performs as well as the GT risk index in maintaining disease control, and can reduce fungicide use even further.

## ACKNOWLEDGEMENTS

We thank A. Albrecht, R. Doody, L. Duffau, J. Enyard, Dr. Zhian Kamvar, T. Neill, T. Nguyen, Dr. F. Peduto Hand, L. Schiller, and Dr. A. Sutherland for technical assistance and discussion and recognize funding from USDA-CRIS project 5358-22000-039-00D,USDA-NIFA project CA-D-PPA-2131-H and the UC Davis Department of Plant Pathology. We also thank Drs. Emerson Del Ponte and David Rizzo as well as three anonymous reviewers for critical reading and useful suggestions of earlier versions of this manuscript. The use of, trade, firm, or corporation names in this publication are for information and convenience of the reader. Such use does not constitute an endorsement or approval by the USDA or the Agricultural Research Service, or by the University of California of any product or service to the exclusion of others that may be suitable.

